# Evolution and remodeling of acceptor specificity in the NodZ α-1,6-fucosyltransferase

**DOI:** 10.64898/2026.10.04.756492

**Authors:** Nicholas T. Smith, George C. diCenzo, Graeme W. Howe

## Abstract

Glycosyltransferases have broad diversity in their acceptor substrates, yet the molecular and evolutionary basis of this diversity remains poorly understood. NodZ is an α-1,6-fucosyltransferase that fucosylates Nod factors, lipochitooligosaccharide molecules involved in the establishment of rhizobium–legume symbioses. While NodZ is the only known bacterial member of the GT23 family of α-1,6-fucosyltransferases, the determinants of its acceptor selectivity and the evolutionary origins thereof are unclear. Here, a phylogenetic analysis of 332 NodZ-like proteins revealed a distinct rhizobial clade and significantly greater structural diversity in the *N*-terminus than in the *C*-terminus. Kinetic analysis of a *Mesorhizobium japonicum* NodZ demonstrated selectivity for longer chitooligosaccharide acceptors (comprised of four to five GlcNAc units), while alanine scanning revealed W139 as a primary determinant of this chain length preference. We propose that W139 forms an aromatic platform that engages distal GlcNAc units present only in longer acceptors, coupling productive binding to acceptor chain length. We then used the NodZ phylogeny to reconstruct ancestral NodZ homolog sequences, which indicated that W139 emerged at the base of the rhizobial clade, and thus may represent a key evolutionary step towards canonical NodZ function. Additionally, the most recent resurrected ancestor was capable of fucosylating chitooligosaccharides with lower efficiency than the extant NodZ, suggesting that efficiency towards chitooligosaccharides was improved during the evolution of this family. Results described here demonstrate that evolutionary diversification of the *N*-terminal region can remodel acceptor specificity while preserving the conserved catalytic architecture of GT23 fucosyltransferases, providing a framework for understanding and engineering fucosyltransferase specificity.

## INTRODUCTION

Fucosyltransferases are of significant interest because of the impact that fucosylation has on several distinct processes, including cancer progression (1), bacterial pathogenesis (2), and plant-microbe signalling (3). While all known fucosyltransferases employ the nucleotide sugar guanosine 5′-diphosphate-β-L-fucose (GDP-Fuc) as the glycosyl donor, there is enormous diversity in their acceptor substrates (4). These enzymes are thus classified in the carbohydrate-active enzymes (CAZy) database (http://www.cazy.org/) (5) according to the nature of the acceptor and the glycosidic linkage formed.

The molecular evolution of fucosyltransferases has resulted in a wide range of enzymes with diverse functions. For instance, the CAZy GT23 family encompasses fucosyltransferases that all form α-1,6 linkages, but that use different acceptors and fulfill different biological roles. The first and most well-studied GT23 member is FUT8, a eukaryotic enzyme that installs α-1,6 linked L-fucose on the asparagine-linked *N*-acetylglucosamine (GlcNAc) unit in *N*-glycans (6). Another GT23 member is NodZ, an enzyme only identified in a polyphyletic group of nitrogen-fixing bacteria known as rhizobia (7). Rhizobia are soil bacteria that can associate with legume plants (e.g., peas and beans); during this association, rhizobia fix atmospheric N_2_ into NH_3_ and provide this bioavailable source of N to the plant (8) in a mutualistic relationship that is highly relevant to agricultural biotechnology. The symbiosis between rhizobia and legumes is initiated following an exchange of signals; plants release flavonoids, and in response, rhizobia produce Nod factors (9). Rhizobial Nod factors are lipochitooligosaccharide (LCO) signalling molecules comprised of three to six β-1,4-linked *N*-acetylglucosamine (GlcNAc) units in which the nonreducing terminal monomer is acylated with a fatty acid (**Figure S1**) (10). In some rhizobia, NodZ catalyzes the formation of an α-1,6 linkage between L-fucose and the reducing terminal GlcNAc of the Nod factor (7). Fucosylation of the Nod factor is required for recognition by some plant receptors, meaning that rhizobia that encode NodZ are able to interact with different plants than those who do not (3, 11).

Due to its role in biosynthesizing the signal that initiates rhizobium-legume symbioses, understanding how NodZ functions could inform future efforts to manipulate host-microbe specificity. Crystallographic analysis of the NodZ from *Bradyrhizobium* sp. WM9 revealed that the enzyme adopts the GT-B fold found in diverse glycosyltransferases (12, 13). This fold is comprised of two Rossmann-like (α, β, α) domains separated by a flexible linker, where the *C*- and *N*-terminal sides of the linker are presumed to be responsible for donor and acceptor binding, respectively (14, 15). In line with this, the structure of the GDP-bound NodZ from *Bradyrhizobium* sp. WM9 revealed seven donor binding residues within the *C*-terminus and only three donor binding residues within the *N*-terminus (12, 13). Mutagenesis experiments using the NodZ from *Azorhizobium caulinodans* ORS571 identified *C*-terminal residues critical for NodZ function, identifying arginine and aspartate residues that are conserved with FUT8, as well as histidine and asparagine residues that are not conserved with FUT8 (16). Moreover, the removal of a predicted α helix made up of the 65 *C*-terminal residues of the NodZ of *Sinorhizobium fredii* NGR234 did not abolish *in vitro* activity, highlighting regions of the NodZ protein that may be dispensable for catalysis (17).

Previous works have provided some insight into how NodZ interacts with its acceptor substrate. The NodZ proteins from *B. japonicum* and *S. fredii*, for instance, were shown to exhibit higher activity with chitooligosaccharide (CO) acceptors like *N*,*N*’,*N*’’,*N*’’’,*N*’’’’-pentaacetyl chitopentaose ((GlcNAc)_5_; CO5; **Figure S1**) than for monomeric GlcNAc or for LCOs (18, 19). This finding led to the hypothesis that NodZ fucosylates CO substrates before the acyl chain is installed on the nonreducing terminus of the Nod factor *in vivo* (18, 19). Although they are not preferred substrates, NodZ also exhibits promiscuity towards various mono- and disaccharide acceptors, allowing the biocatalytic assembly of fucosylated compounds from non-cognate substrates (20). NodZ also exhibits low fucosylation activity on both the branched M3N2 *N*-glycan and the Lewis-X epitope (18), showing that although the enzyme prefers linear GlcNAc oligomers as acceptors, other acceptor substrates can be fucosylated by NodZ.

Since it is the only known bacterial GT23 member, NodZ is frequently compared to the eukaryotic fucosyltransferase, FUT8 (13, 16, 21–25). However, there are significant differences between these two enzymes. FUT8 encodes a type II *N*-terminal transmembrane helix that allows its dimerization (26) and a *C*-terminal Src homology 3 (SH3) domain that is essential for catalysis and subcellular localization to the Golgi apparatus (12, 27), both features that are absent from NodZ. Although FUT8 shows acceptor promiscuity towards COs, FUT8-catalyzed modification of CO5 with GDP-Fuc occurs mainly at C6 of the central GlcNAc unit, not the reducing terminal sugar that NodZ targets (28, 29). This, coupled with the structural similarity between the catalytic domains of FUT8 and NodZ (12, 30), suggests divergence within the regions responsible for positioning the acceptor. While the acceptor-binding regions in FUT8 have been somewhat characterized (6, 21), how NodZ binds its CO substrates to facilitate the installation of the critical L-fucose recognition element during Nod factor biosynthesis remains unclear. As NodZ adopts a GT-B fold, acceptor-specific motifs are likely to be concentrated in the *N*-terminal domain, but no experimental evidence is available to provide further insights into how this domain binds COs, such that its role in acceptor specificity remains entirely uncharacterized.

To track the evolution of NodZ and its CO binding motifs, we assembled 332 putative homologs of this enzyme. Models of these proteins showed more structural variability in their *N*-termini than their *C*-termini, suggesting that diversification of *N*-terminal acceptor-binding regions may have been an important event during the evolutionary history of NodZ. With a combination of *in silico* predictions, site-directed mutagenesis, and kinetic assays, we elucidated four residues that contribute to acceptor recognition and/or catalysis, and we illustrate that the importance of these residues can vary depending on the chain length of the acceptor CO substrate. We then conducted ancestral sequence reconstruction (ASR) to “resurrect” predicted ancestral NodZ proteins, identifying the emergence of key acceptor-binding residues at the node representing the ancestor of all known rhizobial NodZ enzymes. Further characterization of the ancestral proteins revealed that enhancements in catalytic efficiency and mutations in the *N*-terminal domain may have been key to the evolution of function within extant NodZ proteins. Collectively, these results provide a robust understanding of the evolution and function of the α-1,6-fucosyltransferase family that will inform the development of engineered fucosyltransferases with non-native selectivities for use in agriculture and medicine.

## RESULTS AND DISCUSSION

### Rhizobial NodZ proteins are phylogenetically distinct from related fucosyltransferases

To compile a set of proteins related to NodZ, a modified reciprocal hit approach was used: the UniProt database was queried using the Pfam NodZ Hidden Markov Model (HMM) PF05830, then hits were screened against an HMM library to identify those proteins whose best scoring match was PF05830. This method yielded a filtered set of 332 putative NodZ-like proteins. While 74% of these sequences belonged to bacteria, 20% were eukaryotic, and 6% were viral, archaeal, or unclassified. In addition to the diverse organisms of origin, a maximum-likelihood phylogeny of these sequences (**Figure 1A**) contained long branch lengths, indicating sequence dissimilarity amongst the proteins. An exception are the characterized NodZ proteins from rhizobia, which populated a monophyletic clade of 30 proteins with short branches; all 30 proteins in this clade are from rhizobia belonging to the order *Hyphomicrobiales*. The 30 rhizobial NodZ proteins were grouped into one cluster in a sequence similarity network (SSN) built with the same protein set and a clustering cutoff of approximately 25% sequence ID (**Figure 1B, 1C**). This “rhizobial” SSN cluster additionally included 9 proteins from non-rhizobial bacteria, which formed sister lineages to the rhizobial clade in the phylogeny.

**Figure 1.**
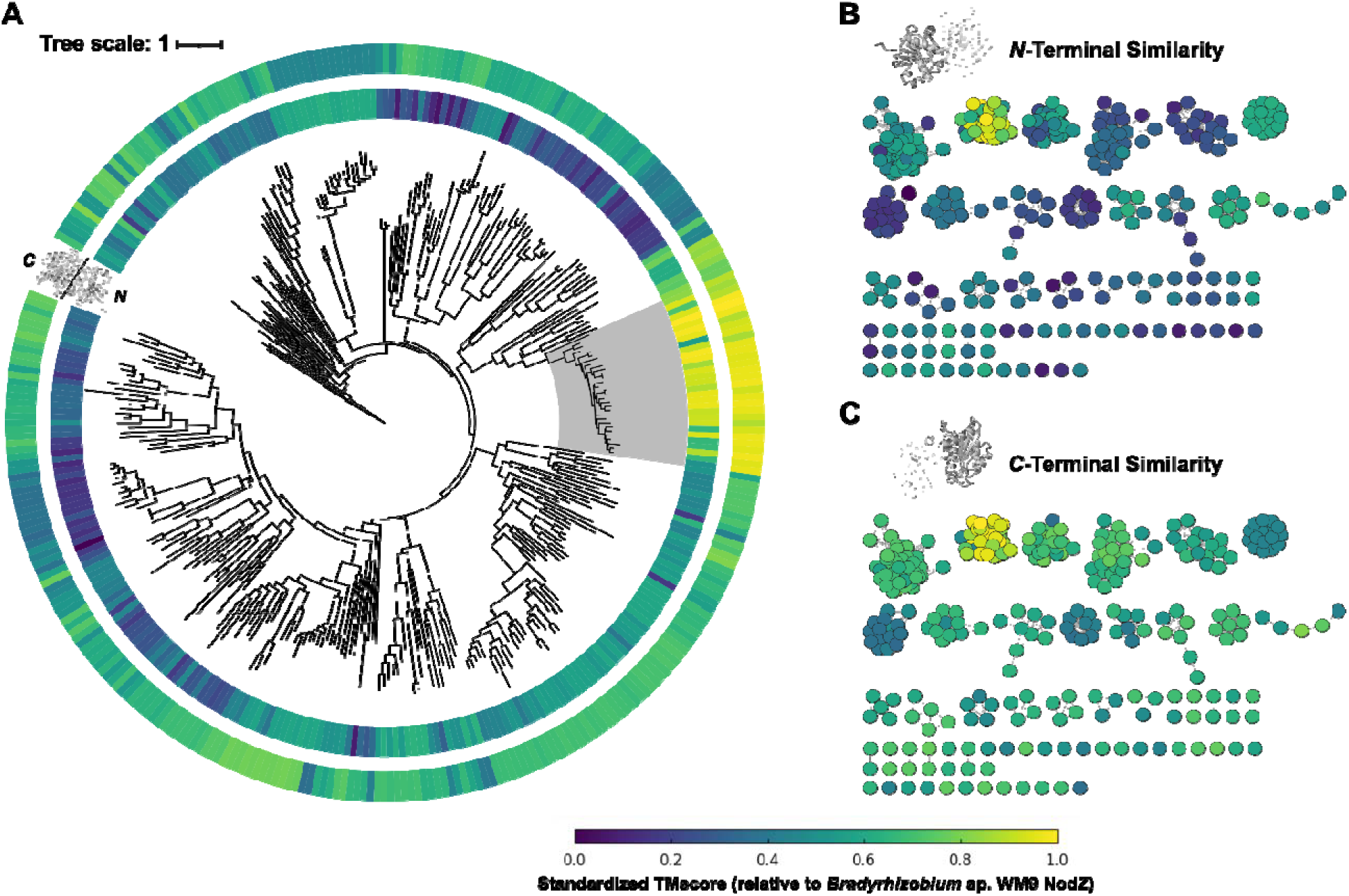
Molecular evolution of putative NodZ-like proteins. The phylogenetic analysis of putative NodZ-like proteins is shown. (***A***) A maximum-likelihood phylogeny generated from 332 NodZ-like protein sequences. The scale bar represents the average number of amino acid substitutions per site. All protein in the phylogeny were modelled with AlphaFold2 and the models were truncated at the linker region separating *N* and *C*-termini, then the truncated structures were aligned to the full-length reference model of *Bradyrhizobium* sp. WM9 NodZ using TM-align. Colours represent TM-align’s TMscore for the alignment of truncated *N* or *C*-terminus structures to the full-length reference model. The inner datastrip represents *N*-terminus homology and the outer represents *C*-terminus homology. The “rhizobial clade” of 30 proteins is shaded in grey. An interactive form of this phylogeny can be accessed at https://itol.embl.de/shared/2CgM1MzDxOeqC. (***B***) A sequence-similarity network generated from the 332 NodZ-like proteins. Nodes (circles) represent protein sequences, with edges (lines) connecting nodes that share ∼25% sequence identity or higher. The nodes are coloured based on predicted *N*-terminus structural homology to the NodZ reference. (***C***) The same sequence similarity network as in panel B, but coloured based on predicted *C*-terminus structural homology to the reference.

The protein set was also scanned for homology to HMMs built for the CAZy (version 13) families (5). As expected, the vast majority (93%) of the proteins were best classified within the GT23 family; however, a small subset belonged to the GT11 (6%) or GT37 (0.3%) families. All three of these CAZy families contain fucosyltransferases, but GT23 enzymes form α-1,6 linkages, whereas GT11 and GT37 encompass a mixture of α-1,2 and α-1,3 fucosyltransferases (5).

Previous work with the NodZ of *A. caulinodans* ORS571 identified specific arginine and aspartate residues as essential for catalysis (R182 and D275 in *A. caulinodans* NodZ) (16), while work with the NodZ of *Bradyrhizobium* sp. WM9 identified specific histidine and serine residues required for binding to GDP (H175 and S287 in *Bradyrhizobium* sp. WM9 NodZ) (13); these four residues were nearly 100% invariant across the 332 NodZ-like proteins (**Table S1**). In contrast, the NodZ-specific histidine and asparagine residues (H183 and N185 in *A. caulinodans* NodZ), whose replacement with the analogous residues from other fucosyltransferases results in a loss of activity (16), were only 19% and 15% conserved, respectively, across the 332 NodZ-like proteins. Other GDP-binding residues from characterized NodZ homologs (13) ranged from 12-88% conservation throughout this dataset.

Taken together, the residue conservation and HMM analyses suggest that the rhizobial NodZ protein sequences are highly conserved with one another, but distinct from similar proteins. They further suggest that the “NodZ-like” proteins comprising the rest of the phylogeny are likely α-1,6-fucosyltransferases, but that they may not possess NodZ activity. These NodZ-like proteins might represent additional GT23 members with unexplored acceptor selectivities and/or biological roles.

### The specialized structure of the rhizobial NodZ N-terminus

Structural conservation for the collected NodZ-like proteins was evaluated using models generated with AlphaFold2 (31). Since the *N*- and *C*-terminal Rossmann folds of GT-B glycosyltransferases generally reflect respective acceptor and donor binding regions (14, 15), we truncated each modelled protein at the linker between the *N-* and *C-*termini. The average confidence (pLDTT) was 87 for the *N*-terminal models and 89 for the *C*-terminal models, indicating that the confidence was not biased towards either domain. Next, we used TM-align (32) to align the models of each domain to the model of NodZ from *Bradyrhizobium* sp. WM9. This enzyme has been characterized structurally but since available crystal structures are missing some segments (12, 13), the AlphaFold2 model was used as the full-length NodZ reference. As expected, the highest structural homology scores were found within the 30-member rhizobial clade (**Figure 1**), indicating that structural conservation amongst NodZ proteins was high. Additionally, across all 332 NodZ-like proteins, 87% aligned to the reference with a higher score for their *C*-terminus than for their *N*-terminus (**Figure 1B, 1C**). This result suggests that there is higher average conservation of the structure of the *C*-terminus relative to that of the *N*-terminus across the 332 NodZ-like proteins. The *C*-terminus of NodZ is predominately involved in GDP-binding, whereas the *N-*terminus is likely to be involved in acceptor recognition (13). Consequently, we interpret the structural similarity data as further support for the hypothesis that the 332 NodZ-like proteins have a shared ability to bind (and likely activate) GDP-linked sugars for glycosylation, but that these putative fucosyltransferases may recognize diverse acceptor molecules due to sequence variation in their *N*-terminal domains.

### Prediction of acceptor-binding motifs in the N-terminal domain

The residues used by NodZ to interact with the GDP moiety of the GDP-Fuc donor substrate are known, but crystals in complex with an acceptor substrate have proven difficult to obtain (13). Therefore, an *in silico* approach was employed to predict the binding site of the canonical NodZ CO substrate, CO5 (18, 19). As a template for this, we used the *M*. *japonicum* R7A NodZ protein (*Mj*NodZ). This protein is known to be functional *in vivo* (3), but no *in vitro* kinetic characterization of NodZ proteins from *Mesorhizobium* have been reported, making *Mj*NodZ an interesting scaffold for deeper characterization. First, the sequence of *Mj*NodZ, along with the ligands GDP-Fuc and CO5, was submitted to the Chai1 server (33) for co-modelling. In the predicted *Mj*NodZ-GDP-Fuc-CO5 complex, the GDP-Fuc donor was positioned in the donor-binding pocket, and several residues predicted to contact this substrate were found in coincident positions within co-crystal structures of diverse fucosyltransferases (**Figure S2)**. The CO5 acceptor substrate was placed in the nearby groove between the *N* and *C*-termini (**Figure 2**). As the reducing terminal sugar of CO5 was oriented reasonably close to the GDP-Fuc substrate, this structural model was taken to provide a reasonable first estimate of the binding pocket for the sugar-acceptor substrate.

**Figure 2.**
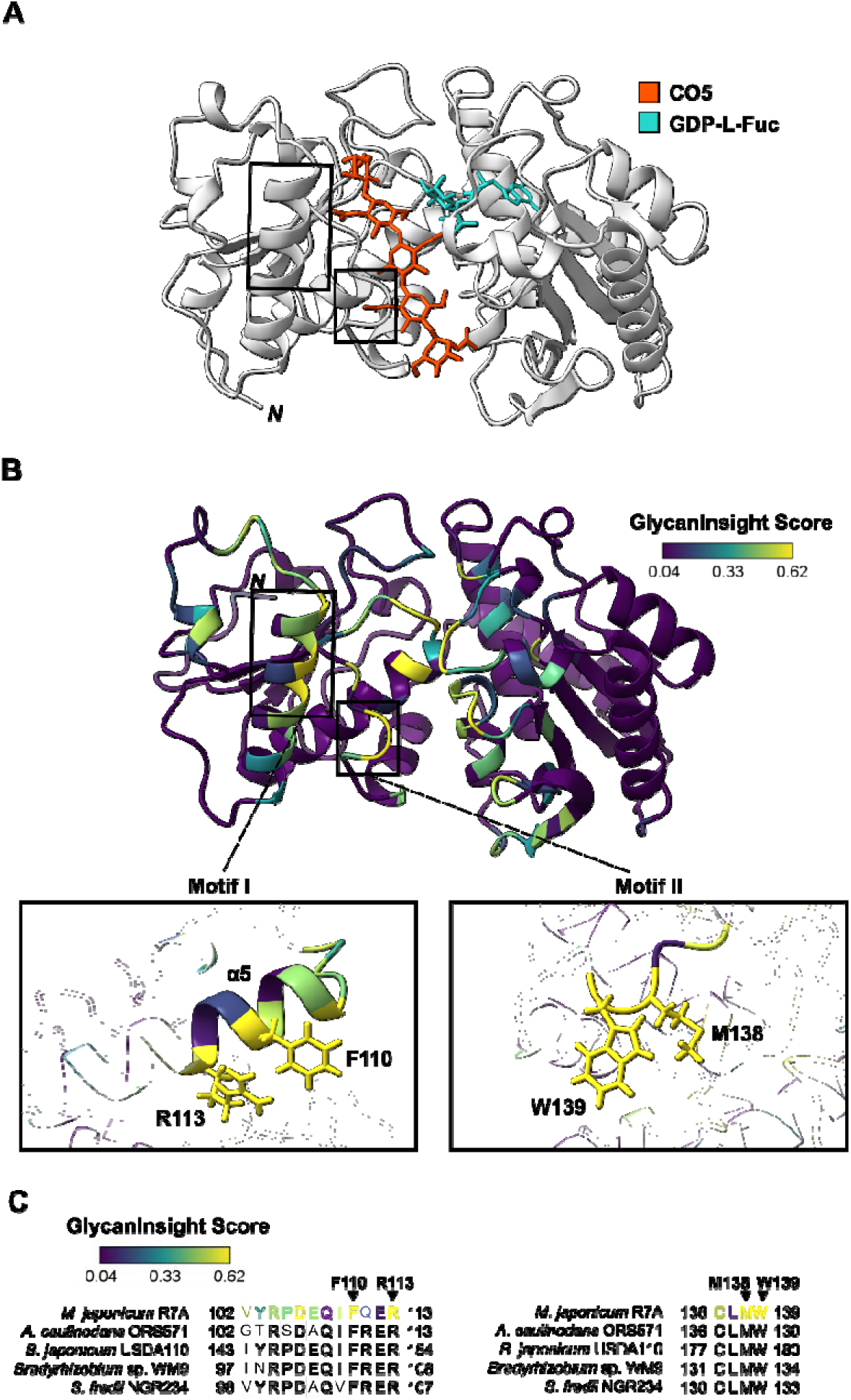
Putative NodZ chitooligosaccharide-binding motifs. The results of an *in silico* prediction of chitooligosaccharide-binding residues in NodZ are presented. (***A***) The Chai1-predicted *Mj*NodZ comple with CO5 (red) and GDP-Fuc (cyan). The program inputs were the amino acid sequence of *Mj*NodZ and SMILES-formatted ligands for CO5 and GDP-Fuc. The regions encoding motifs I and II are shown with boxes. (***B***) The results of a GlycanInsight prediction of CO5-binding residues. The GlycanInsight program inputs were the AlphaFold2 model of *Mj*NodZ and the GLYCAM-WEB optimized CO5 model. The top frame depicts the full *Mj*NodZ AlphaFold2 model with residues coloured based on GlycanInsight score, as indicated in the gradient legend. The regions encoding motifs I and II are shown with boxes. The bottom frame depicts closer views of motifs I (residues 102-113 in *Mj*NodZ) and II (residues 136-139 in *Mj*NodZ). The amino acid side chains are visible only for residues mutated to alanine in this study. (***C***) A multiple sequence alignment showing motifs I and II, and surrounding residues, of functionally annotated NodZ enzymes. The rhizobium of origin for each sequence is indicated; further details on these sequences can be found in **Table S8**. For the *Mj*NodZ sequence, residues are coloured according to their GlycanInsight score. A grey box is used to indicate residues that are 100% conserved across the five proteins. Arrows are used to indicate the residues that were mutated to alanine in this study.

For more quantitative evaluation of this model, an optimized model of the CO5 ligand was generated with the GLYCAM-Web tool (34), and this conformer was submitted with the AlphaFold2 model of apo-*Mj*NodZ to GlycanInsight (35), a machine learning-enabled tool that predicts the likelihood of each residue in a given protein to be involved in sugar binding/recognition (**Table S2**). GlycanInsight predicted likely CO5-binding residues (defined by GlycanInsight scores greater than 0.5) on both sides of the groove between the *N*- and *C*-terminal domains, with the highest probability scores found in the *N*-terminal domain. Notably, the highest-confidence CO5-binding residues were found within regions that lined the predicted CO5-binding groove in the Chai1 *Mj*NodZ-GDP-Fuc-CO5 model (**Figure 2**). These residues were concentrated in three discrete regions: the α1 helix, the α5 helix, and the loop preceding the α6 helix (**Figure S3, Table S3**). The α1 helix encodes two residues (G21 and D22) known to interact with GDP-Fuc in the *Bradyrhizobium* sp. WM9 NodZ crystal structure (13), suggesting a role in donor rather than acceptor binding. As such, efforts to identify determinants of acceptor recognition focused instead on the α5 helix and the pre-α6 helix loop, termed motifs I and II, respectively. To test the hypothesis that these were CO-binding motifs, two conserved NodZ residues were selected per motif and prioritized for biochemical analysis.

### Residues in motifs I and II are critical for CO3, CO4, and CO5 fucosylation

Based on high GlycanInsight scores and conservation with known NodZ enzymes, residues F110 and R113 in motif I, and M138 and W139 in motif II, were targeted for alanine scanning mutagenesis, after which the mutant *Mj*NodZ enzymes were expressed in *Escherichia coli* BL21 (DE3) and purified with immobilized metal affinity chromatography (IMAC) (**Figure S4**). Enzyme activity was then assessed with GDP-Fuc as a donor and a panel of COs of varying length (from 2 to 5 GlcNAc units) as acceptors.

Reactions were performed in the presence of calf intestinal phosphatase (CIP) to catalyze hydrolysis of the GDP released during the reaction and enable initial rate measurements with a malachite green phosphate-detection assay (**Figure S5**).

As expected, wildtype *Mj*NodZ exhibited fucosylation activity toward each tested CO. Notably, Michaelis-Menten kinetics revealed substantially higher *K*_M_ values for CO2 and CO3 than for CO4 and CO5, whereas similar *k*_cat_ values were observed for all four COs (**Table 1, Figure S6**). This indicates that NodZ’s selectivity for CO4 and CO5 is primarily driven by enhanced binding to these substrates rather than increased catalytic turnover. To probe how the identified motifs might be interacting with the CO substrates, initial rates for each alanine mutant were measured with fixed CO concentrations that were >4x the *K*_M_ observed with the wildtype enzyme (**Figure 3A**). Mutations in motif I (F110A and R113A) resulted in severe reductions in activity (<15% of wildtype activity) for CO5, CO4, and CO3, but significantly more activity (∼25-40%) was retained against CO2. The activities of the motif II mutants, M138A and W139A, were more variable. The M138A mutation abolished almost all activity on each CO, while the effect of the W139A mutation was highly substrate-specific, with this mutant losing nearly all activity with CO5 and CO4 but exhibiting ∼30% of wildtype activity with CO3 and ∼90% with CO2. Collectively, these results suggest that the four mutated residues play distinct roles in coordinating COs of different lengths. Whereas all four residues were critical for efficient catalysis with CO3, CO4, and CO5, the residues varied in their importance for catalysis of CO2, with W139 appearing to be required for efficient fucosylation of longer COs (CO4 and CO5) but dispensable for fucosylation of CO2.

**Figure 3.**
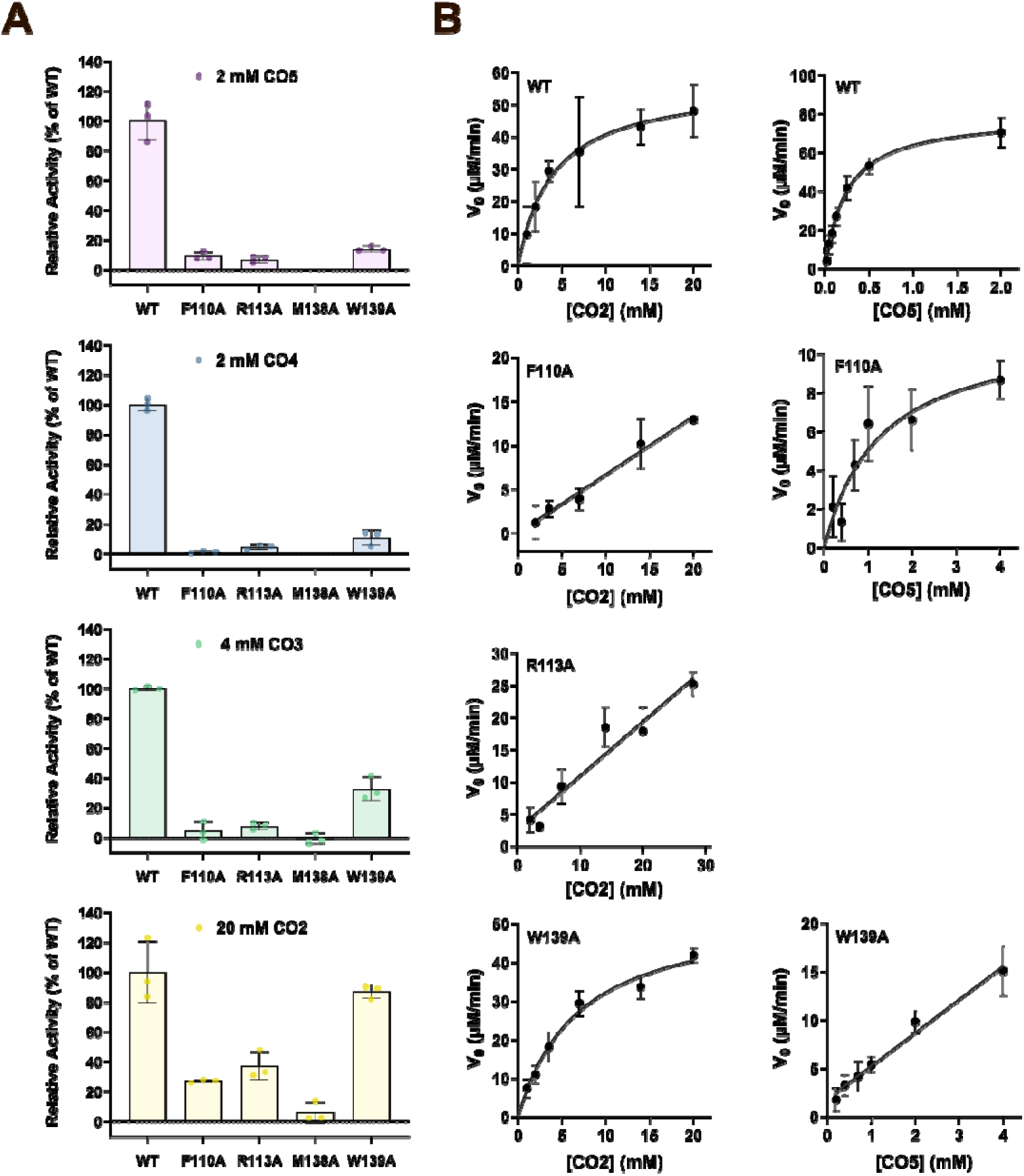
Kinetic effects of NodZ point mutations. The results of *in vitro* fucosyltransferase kineti assays are shown. (***A***) The initial reaction velocities of each *Mj*NodZ point mutant are expressed a percentages relative to the wildtype enzyme (WT), which is set to 100%. Each bar plot represents the results for a different CO acceptor substrate. Individual data points are shown with coloured circles. Error bars represent standard deviation from the mean. Where bars are not shown, a time-dependent linear absorbance change could not be observed at all. (***B***) Plots of initial reaction velocity against substrate concentration are shown for all *Mj*NodZ mutants with sufficiently measurable activity. Each point is the mean of at least three replicates and error bars represent standard deviation from the mean. The data are fitted with the Michaelis-Menten equation, or, if saturation was not observed, linear regression. Data were fit to the relevant equation using GraphPad Prism, and the derived parameters are given in **Table 2**. Data are now shown of M138A or for W139A with CO5 as it was not possible to fit a line or curve to these data.

**Table 1.** Kinetic parameters of *Mj*NodZ with COs of varying lengths.

| Substrate | $K_M$ ($\mu\text{M}$) | $k_{\text{cat}}$ ($\text{s}^{-1}$) | $k_{\text{cat}} / K_M$ ($\text{M}^{-1}\text{s}^{-1}$) |
| --- | --- | --- | --- |
| CO5 | $270 \pm 30$ | $0.42 \pm 0.02$ | $1800 \pm 200$ |
| CO4 | $200 \pm 40$ | $0.44 \pm 0.03$ | $2200 \pm 500$ |
| CO3 | $1500 \pm 500$ | $0.36 \pm 0.04$ | $250 \pm 80$ |
| CO2 | $4000 \pm 500$ | $0.303 \pm 0.016$ | $76 \pm 12$ |

**Table 2.** Kinetic parameters of *Mj*NodZ point mutants with CO5 and CO2.

| | Model Used | $K_M$ ($\mu\text{M}$) | $k_{\text{cat}}$ ($\text{s}^{-1}$) | $k_{\text{cat}}/K_M$ ($\text{M}^{-1}\text{s}^{-1}$) |
| --- | --- | --- | --- | --- |
| **Substrate: CO5** |  |  |  |  |
| *MjNodZ* | Michaelis-Menten | $270 \pm 30$ | $0.42 \pm 0.02$ | $1800 \pm 200$ |
| F110A-*MjNodZ* | Michaelis-Menten | $1300 \pm 500$ | $0.06 \pm 0.01$ | $48 \pm 20$ |
| R113A-*MjNodZ* | N.A.* | N.D.† | N.D.† | N.D.† |
| M138A-*MjNodZ* | N.A.* | N.D.† | N.D.† | N.D.† |
| W139A-*MjNodZ* | Linear | >>WT | N.D.† | $20.1 \pm 1.0$ |
| **Substrate: CO2** |  |  |  |  |
| *MjNodZ* | Michaelis-Menten | $4000 \pm 500$ | $0.303 \pm 0.016$ | $76 \pm 12$ |
| F110A-*MjNodZ* | Linear | >>WT | N.D.† | $3.5 \pm 0.4$ |
| R113A-*MjNodZ* | Linear | >>WT | N.D.† | $4.5 \pm 0.4$ |
| M138A-*MjNodZ* | N.A.* | N.D.† | N.D.† | N.D.† |
| W139A-*MjNodZ* | Michaelis-Menten | $6900 \pm 1200$ | $0.29 \pm 0.02$ | $42 \pm 8$ |

\* N.A. refers to kinetic datasets which could not be fitted to either model, due to insufficient activity.
† N.D. refers to parameters that were not determined due to inability to saturate the enzyme or insufficient activity.

### Motif I and II residues affect NodZ efficiency and CO length preference

To probe the specific roles of the mutated residues more directly, we attempted to use Michaelis-Menten kinetics to determine how the kinetic parameters *k*_cat_ and *K*_M_ changed in response to each mutation. Specifically, we assayed F110A, R113A, and W139A with a range of concentrations of CO2 and CO5, representing the shortest and longest COs used. In many cases, these mutants had significantly impaired substrate utilization, such that saturation could not be observed even with very high CO concentrations (**Figure 3B**). Although this suggested drastic increases in *K*_M_, the linear relations between initial rate and [CO] allowed only for catalytic efficiencies (*k*_cat_/*K*_M_) to be determined where saturation was not achieved.

For other mutant-substrate pairs, activity was too low to allow for reliable measurement, thus precluding the determination of these kinetic parameters. The resulting kinetic measurements are summarized in **Table 2**.

In the predicted structure, the selected motif I residues F110 and R113 were located on a helix near the putative active site (**Figure 2**), so alanine mutants at either position were predicted to disrupt binding to the reducing terminus of the CO acceptors. We observed that the F110A mutation decreased catalytic efficiency (*k*_cat_/ *K*_M_) by ∼37-fold for CO5 and ∼20-fold for CO2 compared to the wildtype enzyme (**Table 2**). For CO5, this could be attributed to an approximate seven-fold decrease in *k*_cat_ and five-fold increase in of *K*_M_, implicating this residue in both turnover and binding with CO5. With up to 20 mM CO2, the F110A mutant showed no signs of saturation, indicating that this mutant has a much higher *K*_M_ for CO2 than the value of *K*_M_ = 4.0 ± 0.5 mM observed with wildtype *Mj*NodZ. Therefore, F110 likely contributes to binding of both CO5 and CO2, although the extent of this effect with CO2 could not be quantified. On the other hand, the R113A mutation disrupted CO5 fucosylation so severely that initial reaction rates with [CO5] of up to 4 mM were too low to distinguish a substrate-dependent response. The R113A mutant had measurable activity with CO2 but still exhibited a ∼17-fold reduction in *k*_cat_/ *K*_M_ with this substrate, relative to the wild type. As with the F110A mutant, R113A could not be saturated with CO2, suggestive of an increase in *K*_M_ that was too large to be quantified with reasonable substrate concentrations. Overall, the dramatic loss of activity and apparent substrate binding in both motif I mutants suggests that F110 and R113 play a vital role in correctly orienting both short and long CO acceptors for the sugar transfer to take place.

The selected residues in motif II (M138 and W139) were predicted to form a loop that interacts with the nonreducing terminal of CO5 (**Figure 2**). Surprisingly, initial kinetic analysis suggested that M138A lost nearly all detectable activity with each of the substrates tested. In the *Bradyrhizobium* sp. WM9 NodZ-GDP crystal structure (13), the sulfur of the analogous Met residue (M133) protrudes into the cleft between the *N-* and *C*-terminal domains. The M138 sulfur might interact with W139 (or other nearby aromatic residues) via S-π interactions that modulate the positioning and/or electronics of the aromatic side chain. While more experimental evidence would be required to further probe the role of M138 in *Mj*NodZ, a second-shell interaction of the sort proposed would be consistent with the recognized importance of CH-π interactions in glycan recognition (36), as well as the role of aromatic residues in some human fucosyltransferases in acceptor substrate selectivity (37). Indeed, mutation of the aromatic W139 reduced *k*_cat_/*K*_M_ for CO5 ∼90-fold compared to the wild type (**Table 2**). One contributing factor was a dramatically increased *K*_M_ for CO5, since W139A was not saturated at [CO5] >14x the *K*_M_ of the wildtype enzyme (**Figure 3B**). This indicates that the W139A mutation severely reduced binding to CO5. In striking contrast, W139A was minimally impaired in its ability to fucosylate CO2, as its *k*_cat_/*K*_M_ value for CO2 was only reduced by ∼1.8-fold relative to that of the wild type. W139A could be saturated with CO2, yielding *K*_M_ and *k*_cat_ values of similar magnitude as the wild type. This substrate-specific phenotype was so pronounced that the apparent catalytic efficiency of the W139A mutant was lower for the “native” substrate, CO5, than for the shorter CO2 (**Figure 3B**). Moreover, W139A was the only mutant that had an incremental activity profile in which the initial rate (expressed as relative activity compared to the wildtype enzyme) was decreased with each additional GlcNAc unit on the acceptor (**Figure 3A**). Consistent with our *in silico* model for CO5 binding by *Mj*NodZ, we propose that CO2 does not extend far enough through the acceptor binding groove to permit W139 to contact this substrate, but with longer CO substrates harboring additional GlcNAc units, the interaction with W139 becomes increasingly important for fucosylation, in a CO-length specific manner.

### The N-terminus of ancestral NodZ proteins encodes additional determinants of function

Although the discovery of functional *N*-terminal motifs provides insight into how NodZ recognizes its acceptor, we also wanted to understand how these regions evolved. To this end, we located a common ancestor (“N0”) of the rhizobial clade and related proteins, then generated a phylogeny with all 48 of its descendants (**Figure 4A**). To predict the ancestral proteins that gave rise to these extant proteins, we used Graphical Representation of Ancestral Proteins (GRASP) (38) to perform ancestral sequence reconstruction (ASR). This resulted in the prediction of six ancestors of the rhizobial NodZ clade, which shared between 38 % and 75 % sequence identity with *Mj*NodZ (**Table S4**).

**Figure 4.**
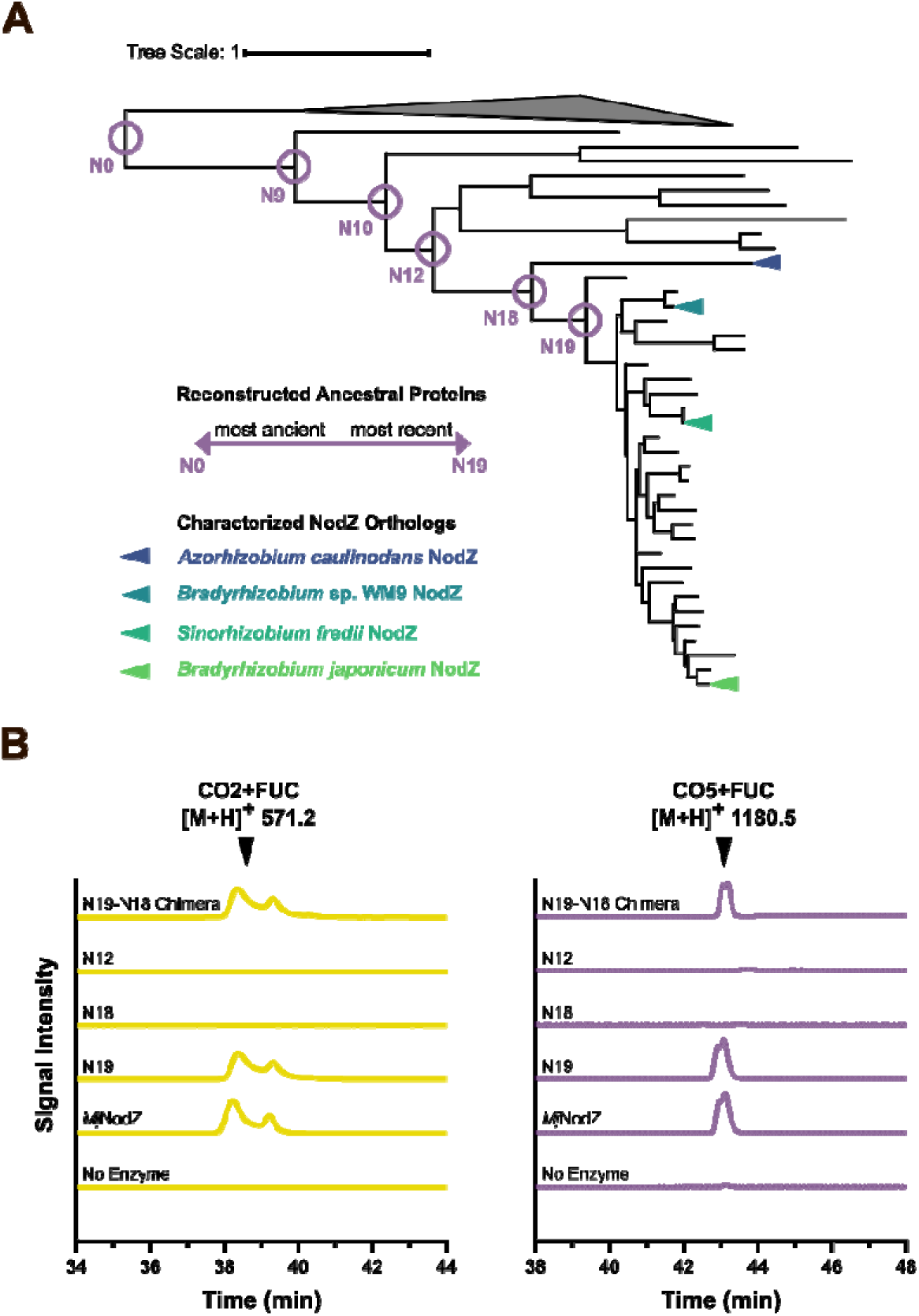
*N*-terminal determinants of activity in ancestral NodZ proteins. The analyses used to reconstruct and characterize ancestral NodZ proteins are presented. (***A***) A maximum-likelihood phylogeny of the rhizobial clade of NodZ and NodZ-like proteins, rooted with an outgroup of NodZ-like algal proteins (shown as collapsed). The rhizobial clade of 30 proteins is shaded in grey. The nodes from which sequences were reconstructed in this study are circled. Experimentally characterized NodZ enzymes are indicated with coloured arrows. The scale bar represents the average number of amino acid substitutions per site. An interactive form of this phylogeny can be accessed at https://itol.embl.de/shared/2CgM1MzDxOeqC. (***B***) The detection of CO fucosylation in reaction performed with ancestral NodZ enzymes and their mutants is shown. Chromatograms show the results of single-ion monitoring (SIM) for the [M+H]^+^ ion corresponding to the fucosylated CO products.

AlphaFold2 models of the two most “recent” of the reconstructed ancestors, N19 and N18, suggested that these proteins adopt a very similar structure to that of the *Bradyrhizobium* sp. WM9 NodZ, while the predicted structures of the more ancient proteins (N12, N10, N9, and N0) were increasingly dissimilar (**Figure S7**). The ancestors N12, N10, N9, and N0 were more well-conserved with NodZ in their *C*-terminal lobes than in their *N*-terminal lobes, supporting the hypothesis that structural divergence accumulated in the *N*-terminus during NodZ evolution. However, the conservation of the residues that were experimentally characterized (F110, R113, M138, and W139) was found to be restricted only to the descendants of N18 (**Table S3**). In the “older” ancestors (N12 and earlier), only R113 was conserved out of these four residues, suggesting that they are unlikely to be active on COs. In other words, the emergence of the functional motifs identified in this study as required for fucosylation of COs (particularly longer-chain COs) coincides with the evolution of the rhizobial NodZ clade, further supporting the hypothesis that the canonical NodZ function arose through evolution of the *N*-terminal domain. Considering that W139 was implicated in the specific recognition of 5-unit COs, changes in motif II might have been a critical step towards the ability of NodZ to fucosylate COs of the appropriate length for a role in Nod factor biosynthesis.

To get empirical insight into the ancestral reconstruction, the ancestral proteins N12, N18, and N19 were expressed in *E*. *coli* and purified with IMAC (**Figure S8**). The ancestral enzymes were incubated with GDP-Fuc and either the CO5 or CO2 acceptor at 37 °C for approximately 20 hours. Reaction products were detected with hydrophilic interaction liquid chromatography coupled with mass spectrometry (HILIC-MS) (39, 40). An enzyme-free negative control confirmed that [M+H]^+^ ions corresponding to unmodified CO5 and CO2 could be detected (**Figure S9**), while the wildtype *Mj*NodZ reactions produced additional signals consistent with fucosylated versions of these acceptors (**Figure 4B**) that correlated with depletion of the substrates. The ancestral protein N19 could fucosylate CO5 and CO2, but with a *k*_cat_/*K*_M_ of 300 ± 100 M^-1^ s^-1^ for CO5 and 40 ± 20 M^-1^ s^-1^ for CO2 (**Figure S10**), corresponding to catalytic efficiencies ∼6-fold and ∼2-fold lower than the corresponding values for *Mj*NodZ (**Table 1**). The lower catalytic efficiency of N19, particularly towards CO5, compared to *Mj*NodZ may reflect enhancements in catalytic efficiency delivered by natural evolution of these enzymes. On the other hand, neither N18 nor N12 could fucosylate CO5 or CO2. However, fusing the *N*-terminal domain of N19 with the *C*-terminal domain of N18 yielded an active enzyme that could fucosylate both of these acceptors. Overall, these results reveal that the *N*-terminal domain plays a functional role in the transition between inactive and active ancestral NodZ proteins, further supporting the hypothesis that diversification of the *N*-terminal domain contributed to the evolution of canonical NodZ function.

## CONCLUSION

Within the GT23 family of α-1,6-fucosyltransferases, NodZ proteins are poorly understood compared to their eukaryotic counterpart, FUT8. As NodZ proteins play a biological role in agriculturally relevant host-microbe signaling, a deeper characterization of this family would not only refine CAZy annotations, but provide the crucial insight required for the manipulation of fucosylated signaling molecules. Here, we show that rhizobial NodZ proteins form a distinct sub-clade within a broader group of 332 predicted “NodZ-like” fucosyltransferases. High-throughput modelling of all 332 proteins revealed greater structural divergence between the *N*-terminal domains relative to the *C*-terminal domains, suggesting that the *N*-terminal domain has been diversified through evolution to accommodate different acceptor substrates. Thus, the true NodZ proteins (that is, α-1,6-fucosyltransferases that act on rhizobial Nod factors) may be a relatively specific sub-family of a broader “NodZ-like” clade, all of which are distinct from eukaryotic FUT8 proteins. Although not explored in this study, these distantly related NodZ-like proteins may form families of GT23 members with previously undiscovered activities.

As the phylogenetic analyses suggested that NodZ might be relatively specialized for its glycosyl acceptor substrates (i.e., COs), we used Michaelis-Menten kinetics to probe acceptor recognition in recombinantly expressed *M. japonicum* R7A NodZ (*Mj*NodZ). *Mj*NodZ exhibited significantly lower *K*_M_ values for longer COs (CO4 and CO5) compared to the shorter ones (CO3 or CO2), indicating that substrate affinity is a strong driver of NodZ acceptor specificity. We then used structure-based prediction tools to highlight candidate residues contributing to CO acceptor binding. Mutagenesis of four selected residues demonstrated that these residues strongly influence CO fucosylation, depending on the length of the CO acceptor. In particular, the W139A mutation strongly impaired CO5 binding but not CO2 binding, demonstrating that this single residue influences acceptor chain-length specificity. Since ancestral sequence reconstruction predicted that W139 emerged at the base of the rhizobial clade, we propose that this residue was a key mutation that allowed efficient fucosylation of longer COs, enabling a role in Nod factor biosynthesis.

Together, this work refines the evolutionary history of a neglected member of the GT23 family and illuminates a model in which the conserved *C*-terminal domain maintains the core catalytic architecture of NodZ-like fucosyltransferases, while diversification of the *N*-terminal domain enables distinct acceptor specificities. In this study, the combination of widely available *in silico* prediction tools with foundational biochemistry not only served as a powerful hypothesis generation and testing workflow but also built the foundation for future enzyme engineering efforts. The identification of residues (e.g., W139) that influence acceptor chain length provides potential targets for engineering fucosyltransferases with customized acceptor preferences, while the successful domain-swapping experiment suggests that domain-level approaches may be an additional strategy for modifying GT-B glycosyltransferase specificity.

## METHODS

### Phylogenetic analysis of NodZ and related proteins

To locate putative NodZ proteins, we used HMMer version 3.3 to search the UniRef50 database for matches to the Pfam HMM for NodZ (PF05830). The resulting proteins were scanned against an HMM database containing models from Pfam-A version 36.0 (13), TIGRFAMs version 15.0 (14), COGs (15) and Protein Clusters (PRK) databases of NCBI’s Conserved Domain Database (CDD). Only proteins that scored highest with PF05830 were included for our analysis. Proteins below 200 and above 480 amino acids in length, and sequences annotated as fragments, were excluded from the dataset. The UniProtKB accessions corresponding to each UniRef50 cluster were downloaded from UniProt (release 2024_02), and CD-HIT version 4.8.1 (41) was used to cluster proteins with 85 % sequence identity or higher. Four functionally annotated rhizobial NodZ proteins (UniProt: Q43966, Q9AQ17, Q45271, and P55355) and the NodZ protein from *M*. *japonicum* MAFF303099 were re-added to the dataset. The protein sequences were aligned with MAFFT version 7.471 (42) and the alignment was tested against several substitution models (LG, WAG, JTT, Q.pfam, JTTDCMut, DCMut, VT, PMB, Blosum62, Dayhoff) with IQ-TREE version 2.2.2.4 (43, 44). A maximum-likelihood phylogeny was generated using the best model (Q.pfam+R9) and 1000 ultrafast bootstrap replicates. The phylogeny was visualized using the online iTOL webserver (45).

### Domain-specific structural analysis of NodZ and related proteins

To compare the conservation of *N-* and *C*-termini across putative NodZ-like proteins, we modelled each protein using AlphaFold2 (31). Each model was split at the region encoded by position 1052 in the alignment, representing the approximate midpoint of the conserved linker region. This resulted in two PDB files per protein model; one for the *N*-terminus and one for the *C*-terminus. The *N-* and *C*-termini models were then aligned to the model of NodZ from *Bradyrhizobium* sp. WM9 or *Mj*NodZ using TM-align (32). The resulting TMscores were normalized to the length of the query.

### In silico modeling and ligand-binding predictions of MjNodZ

To predict the structure of the *Mj*NodZ-substrate complex, the amino acid sequence of the protein (UniProt ID: A0ABX6MZR4) and SMILES-formatted ligands CO5 and GDP-Fuc were submitted to the Chai1 server (33). To obtain a quantitative prediction of the likelihood of the residues of *Mj*NodZ to be involved in CO5-binding, the optimal structure of the ligand was predicted using GLYCAM-Web (46) and submitted along with an AlphaFold2 model of *Mj*NodZ to the GlycanInsight server, with default parameters, to compute per-residue substrate binding likelihood scores with the DeepGlycanSite algorithm (35).

### Ancestral sequence reconstruction

The amino acid sequences of 48 descendants of a common ancestor (“N0”) of the rhizobial clade were assembled. This refined list of sequences was aligned and tested against substitution models as described above. The best model (JTT+R5) was used to generate a maximum-likelihood phylogeny. The multiple sequence alignment and corresponding phylogeny were submitted to GRASP version 2020.05.05 (38) server for a marginal reconstruction using the JTT option.

### Cloning and site-directed mutagenesis

To perform site-directed mutagenesis, inverse PCR was performed using a pET-28a-*Mj*NodZ plasmid (pGQ0146; **Table S5**) as a template. Primers (**Table S6**) with 5 phosphate groups (Thermo Fisher) were used to induce the desired point mutations. PCR amplicons were digested with DpnI (New England Biolabs) to remove methylated DNA and circularized using T4 ligase (New England Biolabs). To generate the N19-N18 chimera, primers with complimentary overhangs were used to amplify the nucleotides encoding residues 1-152 of N19 and the nucleotides encoding residues 153-329 of N18. The resulting amplicons were assembled into a pET-28a vector that was pre-digested with *Nde*I and *Xho*I using the New England Biolabs HiFi mix (New England Biolabs). All constructs and mutants were confirmed using whole plasmid Oxford Nanopore Technologies sequencing at Plasmidsaurus (Louisville, KY, USA).

### Protein expression and purification

Protein expression plasmids (**Table S5**) were transformed into *E*. *coli* BL21 (DE3). Protein overexpression and purification was performed as described previously (47) with minor alterations. All purification steps were performed at 4 °C. Briefly, one-litre *E. coli* expression cultures were induced with 1 mM IPTG, grown at 16 °C for ∼20 hours, then lysed in lysis buffer (20 mM Tris-HCl, 100 mM NaCl, 10% glycerol (v/v), pH 7.6) with 1 mg/mL lysozyme in a French Press (Sigma Aldrich). Clarified supernatant was incubated with 5 mL His60 Ni Superflow resin (Takara Bio) in a Bio-Rad Econo-Column, before washing the resin with wash buffer (20 mM Tris-HCl, 100 mM NaCl, 20 mM imidazole, 10% glycerol (v/v), pH 7.6) and eluting protein with elution buffer (20 mM Tris-HCl, 100 mM NaCl, 500 mM imidazole, 10% glycerol (v/v), pH 7.6). The samples were concentrated with MWCO 30 kDa Amicon Ultra centrifugal filter units (Sigma) and buffer exchanged into storage buffer (40 mM Tris-HCl, 100 mM NaCl, 10% glycerol (v/v), pH 7.6) with PD-10 columns (Cytiva). Protein samples were analyzed for purity with SDS-PAGE (**Figure S4, S7**) and concentrations were determined using the Bradford assay with bovine γ-globulin as the standard.

### Fucosyltransferase kinetic assays

To monitor the kinetics of the fucosyltransferase reaction, assays were performed in glycosyltransferase reaction buffer (50 mM Tris, 100 mM NaCl, pH 7.5) with 15 U/mL calf-intestinal phosphatase (Quick CIP; New England Biolabs). The concentration of Quick CIP was optimized so as to confirm that it was not limiting the apparent rate of the NodZ-catalyzed reaction. The GDP produced during the NodZ-catalyzed sugar transfer was rapidly hydrolyzed by Quick CIP and the inorganic phosphate (P_i_) product was detected with malachite green reagent (Sigma Aldrich). To measure affinity for various acceptor substrates, the concentration of GDP-L-fucose (Ambeed) was held at 100 µM, which was determined to be a saturating concentration for wildtype *Mj*NodZ and N19. Reactions were initiated with the addition of 3.1 µM *Mj*NodZ or variants thereof (6.3 µM, in the case of N19) and incubated at 37 °C. Aliquots of 55 µL were removed during the phase in which the malachite green-derived absorbance was linear with respect to time (within the first 60 seconds). Each NodZ protein/mutant was tested with the maximum CO substrate concentration used to ensure that the absorbance change was linear with respect to time. Reaction aliquots were quenched by immediate addition to 45 µL malachite green reagent. Colour development was allowed to proceed for 20 minutes, then the absorbance at 620 nm was measured using an Epoch2 microplate reader (BioTek). P_i_ concentrations were then quantified using a P_i_ calibration curve.

### HILIC-MS analysis

To detect NodZ reaction products, reactions consisting of 10 mM GDP-Fuc, 6 mM CO (either CO2 or CO5), and 4 µM NodZ, in 50 mM HEPES (pH 7.4) were incubated at 37 °C for approximately 20 hours. Reactions were quenched by centrifugation with a 10 kDa MWCO filter (VWR) and analyzed using HILIC-MS (39, 40). Analytical HILIC-MS was performed on an Agilent InfinityLab LC/MSD series with a Waters XBridge BEH Amide column, 5 μm, 4.6 x 250 mm at a flow rate of 2.0 mL/min, injection volume of 20 μL (2 mg/ml), with 1% of the flow diverted to the ESI-MS detector using a splitter. Mobile phase A was 10 mM ammonium formate in water, adjusted to pH 4.5 with formic acid; mobile phase B was 10 mM ammonium formate in 90% acetonitrile, adjusted to pH 4.5 with formic acid. The gradient used is detailed in **Table S7**. Single-ion monitoring (SIM) settings of *m/z* = 571.2 or 1180.5 were used to detect the [M+H]^+^ ions corresponding to monofucosylated CO2 or CO5, respectively. Single-ion monitoring (SIM) settings of *m/z* = 425.2 or 1034.4 were used to detect the [M+H]^+^ ions corresponding to unmodified CO2 or CO5, respectively. For all chromatograms, retention times were standardized to the retention time of HEPES.

## DATA AVAILABILITY

Scripts required to replicate the analyses described here are available on GitHub at <u>nsmithbiology/nodz_analyses</u>. Interactive versions of the phylogenetic trees can be accessed on iTOL at https://itol.embl.de/shared/2CgM1MzDxOeqC.

## ACKNOWLEDGEMENTS

We thank Kirsten H. Benidickson for assisting with protein quantification and Dr. Chantelle J. Capicciotti for insightful discussion surrounding glycosyltransferase biochemistry and use of her HILIC-MS instrument.

