## Supplemental Information for "Evolution and remodeling of acceptor specificity in the NodZ α-1,6-fucosyltransferase"

**SUPPORTING INFORMATION FOR: Evolution and remodeling of acceptor specificity in the NodZ α-1,6-fucosyltransferase**

**This file includes:**

Tables S1 to S8 (pages S2 to S9)

Figures S1 to S10 (pages S10 to S19)

References for the Supporting Information (page S20)

**Table S1.** Conservation of experimentally validated residues in NodZ-like proteins.

| **Residue** | **Position in *Mj*NodZ** | **Position in *Bradyrhizobium* sp. WM9 NodZ** | **Position in *Ac*NodZ** | **% Conservation in NodZ-like proteins (n=332)** | **Function** |
| --- | --- | --- | --- | --- | --- |
| G | 21 | 16 | 21 | 49 | Binds GDP (1) |
| D | 22 | 17 | 22 | 22 | Binds GDP (1) |
| Y | 50 | 45 | 50 | 60 | Binds GDP (1) |
| H | 180 | 175 | 180 | 94 | Binds GDP (1) |
| R | 182 | 177 | 182 | 99 | Critical for catalysis (2), binds GDP (1) |
| H | 183 | 178 | 183 | 19 | Contributes to catalysis, but not substrate binding (2) |
| N | 185 | 180 | 185 | 15 | Contributes to catalysis (2) |
| D | 229 | 224 | 229 | 88 | Binds GDP (1) |
| D or E | 275 (E) | 270 (D) | 275 (D) | 96 | Critical for catalysis (2), binds GDP (1) |
| S | 292 | 287 | 292 | 97 | Binds GDP (1) |
| A | 293 | 288 | 293 | 12 | Binds GDP (1) |
| F | 294 | 289 | 294 | 60 | Binds GDP (1) |

**Table S2.** High-confidence GlycanInsight scores for *Mj*NodZ and CO5.

| **Residue (*Mj*NodZ)** | **GlycanInsight Score** | **Domain** |
| --- | --- | --- |
| T18 | 0.565 | *N*-terminal |
| G19 | 0.61 | *N*-terminal |
| D22 | 0.587 | *N*-terminal |
| V102 | 0.521 | *N*-terminal |
| D106 | 0.595 | *N*-terminal |
| F110 | 0.619 | *N*-terminal |
| R113 | 0.608 | *N*-terminal |
| C136 | 0.542 | *N*-terminal |
| M138 | 0.625 | *N*-terminal |
| W139 | 0.62 | *N*-terminal |
| R182 | 0.523 | *C*-terminal |
| I189 | 0.575 | *C*-terminal |
| M190 | 0.573 | *C*-terminal |
| H260 | 0.534 | *C*-terminal |
| T291 | 0.597 | *C*-terminal |
| S292 | 0.525 | *C*-terminal |
| A293 | 0.503 | *C*-terminal |
| L310 | 0.524 | *C*-terminal |

**Table S3.** Putative *N*-terminal carbohydrate-binding residues.

| **Residue (*Mj*NodZ)** | **Predicted Location** | **% Conservation in rhizobial clade (n=30)** | **Putative Motif** |
| --- | --- | --- | --- |
| T18 | Pre-α1 loop | 96.7 | N/A |
| G19 | α1 helix | 100 | N/A |
| D22 | α1 helix | 100 | N/A |
| V102 | Pre-α5 loop | 73.3 | I |
| D106 | α5 helix | 100 | I |
| F110* | α5 helix | 100 | I |
| R113* | α5 helix | 100 | I |
| C136 | Pre- α6 loop | 100 | II |
| M138* | Pre- α6 loop | 100 | II |
| W139* | Pre- α6 loop | 100 | II |

*Residue was mutated to alanine in this study

**Table S4.** Percent identity matrix of ancestral NodZ proteins.

|  | **N0** | **N9** | ***Mj*NodZ** | **N18** | **N19** | **N10** | **N12** |
| --- | --- | --- | --- | --- | --- | --- | --- |
| **N0** | 100 | 61.11 | 38.11 | 45.43 | 43.29 | 50.88 | 48.28 |
| **N9** | 61.11 | 100 | 54.88 | 67.68 | 64.33 | 77.64 | 71.98 |
| ***Mj*NodZ** | 38.11 | 54.88 | 100 | 74.16 | 75.08 | 61.70 | 62.92 |
| **N18** | 45.43 | 67.68 | 74.16 | 100 | 95.14 | 79.03 | 82.07 |
| **N19** | 43.29 | 64.33 | 75.08 | 95.14 | 100 | 75.08 | 78.12 |
| **N10** | 50.88 | 77.64 | 61.70 | 79.03 | 75.08 | 100 | 92.05 |
| **N12** | 48.28 | 71.98 | 62.92 | 82.07 | 78.12 | 92.05 | 100 |

**Table S5.** Plasmids and bacterial strains used in this study.

| **Name** | **Characteristics** | **Reference** |
| --- | --- | --- |
| ***Escherichia coli*** |  |  |
| BL21 (DE3) | *fhuA2 [lon] ompT gal* (*λ DE3) [dcm]* ∆*hsdS* | New England Biolabs |
| DH5α | *fhuA2::IS2 Δ(mmuP-mhpD)169 ΔphoA8 glnX44 ϕ80d[ΔlacZ58(M15)] rfbD1 gyrA96 luxS11 recA1 endA1 rphWT thiE1 hsdR17* | New England Biolabs |
| EcGQ0221 | BL21(DE3) (pGQ0146); Km^R^ | This work |
| EcGQ0258 | BL21(DE3) (pGQ0176); Km^R^ | This work |
| EcGQ0259 | BL21(DE3) (pGQ0177); Km^R^ | This work |
| EcGQ0260 | BL21(DE3) (pGQ0178); Km^R^ | This work |
| EcGQ0272 | BL21(DE3) (pGQ0183); Km^R^ | This work |
| EcGQ0281 | BL21(DE3) (pGQ0188); Km^R^ | This work |
| EcGQ0282 | BL21(DE3) (pGQ0189); Km^R^ | This work |
| EcGQ0283 | BL21(DE3) (pGQ0190); Km^R^ | This work |
| EcGQ0284 | BL21(DE3) (pGQ0191); Km^R^ | This work |
| **Plasmids** |  |  |
| pET-28a | A protein overexpression vector; Km^R^ | Twist Biosciences |
| pGQ0146 | pET-28a vector with gene encoding *Mesorhizobium japonicum* R7A NodZ; Km^R^ | Twist Biosciences |
| pGQ0176 | pET-28a vector with gene encoding ancestrally reconstructed NodZ "N19"; Km^R^ | Twist Biosciences |
| pGQ0177 | pET-28a vector with gene encoding ancestrally reconstructed NodZ "N18"; Km^R^ | Twist Biosciences |
| pGQ0178 | pET-28a vector with gene encoding ancestrally reconstructed NodZ "N12"; Km^R^ | Twist Biosciences |
| pGQ0183 | pET-28a vector with gene encoding residues 1-152 of N19 fused to residues 153-329 of N18; Km^R^ | This work |
| pGQ0188 | pET-28a vector containing *Mesorhizobium japonicum* R7A NodZ gene with F110A mutation; Km^R^ | This work |
| pGQ0189 | pET-28a vector containing *Mesorhizobium japonicum* R7A NodZ gene with R113A mutation; Km^R^ | This work |
| pGQ0190 | pET-28a vector containing *Mesorhizobium japonicum* R7A NodZ gene with M138A mutation; Km^R^ | This work |
| pGQ0191 | pET-28a vector containing *Mesorhizobium japonicum* R7A NodZ gene with W139A mutation; Km^R^ | This work |

Km – kanamycin

**Table S6.** Oligonucleotide primers used in this study.

| **Name** | **Primer Sequence (5’-3’)** | **Usage** |
| --- | --- | --- |
| NS014 | **CTGGTGCCGCGCGGCAGCCA**AATGTTAGTGGATGGAAGTATGC | Forward primer for the N-terminal domain of N19, containing an overlap to pET-28a at the NdeI position |
| NS015 | **ATCTCAGGACGAGGGCGCAG**CTGACGGAAGATTTGCCG | Reverse primer for the N-terminal domain of N19, containing an overlap to the C-terminal domain of N18 |
| NS016 | CTGCGCCCTCGTCCTGAGATTCG | Forward primer for the C-terminal domain of N18 |
| NS017 | **AGTGGTGGTGGTGGTGGTGC**CTACGACGCTGAACAATTCTCTGACTG | Reverse primer for the C-terminal domain of N18, containing an overlap to pET-28a at the XhoI position |
| NS022 | AGGAACGCGATGAACTGAACG | Forward primer to induce F110A mutation in MjNodZ |
| NS023 | GA**GC**AATCTGTTCATCCGGG | Reverse primer to induce F110A mutation in MjNodZ |
| NS024 | AGGAA**GC**CGATGAACTGAACG | Forward primer to induce R113A mutation in MjNodZ |
| NS025 | GAAAAATCTGTTCATCCGGGC | Reverse primer to induce R113A mutation in MjNodZ |
| NS027 | TGCCTGATG**GC**GCGCTGC | Forward primer to induce W139A mutation in MjNodZ |
| NS028 | CGCATCGCACACCACGGTGTTC | Reverse primer to induce W139A mutation in MjNodZ |
| NS029 | ATGCGTGCCTG**GC**GTGGCGC | Forward primer to induce M138A mutation in MjNodZ |
| NS030 | CGCACACCACGGTGTTCGCTTCGC | Reverse primer to induce M138A mutation in MjNodZ |

*Bolded and underlined nucleotides indicate regions not homologous to the template. Primers were synthesized with 5’ phosphate groups.

**Table S7.** Linear gradient for HILIC-HPLC-MS.

| **Time (min)** | **A (%)** | **B (%)** |
| --- | --- | --- |
| 0 | 0 | 100 |
| 10 | 0 | 100 |
| 35 | 10 | 90 |
| 45 | 50 | 50 |
| 50 | 50 | 50 |
| 55 | 0 | 100 |
| 60 | 0 | 100 |

* Mobile phase A was 10 mM ammonium formate in water, adjusted to pH 4.5 with formic acid; mobile phase B was 10 mM ammonium formate in 90% acetonitrile, adjusted to pH 4.5 with formic acid.

**Table S8.** Experimentally characterized NodZ enzymes.

| **Species of origin** | **UniProt Accession** | **Reference** |
| --- | --- | --- |
| *Azorhizobium caulinodans* ORS571 | Q43966 | (2, 3) |
| *Bradyrhizobium* sp. WM9 | Q9AQ17 | (1, 4) |
| *Bradyrhizobium japonicum* USDA110 | Q45271 | (5–8) |
| *Mesorhizobium japonicum* R7A | A0ABX6MZR4 | (9), This work |
| *Sinorhizobium fredii* NGR234 | P55355 | (10–12) |

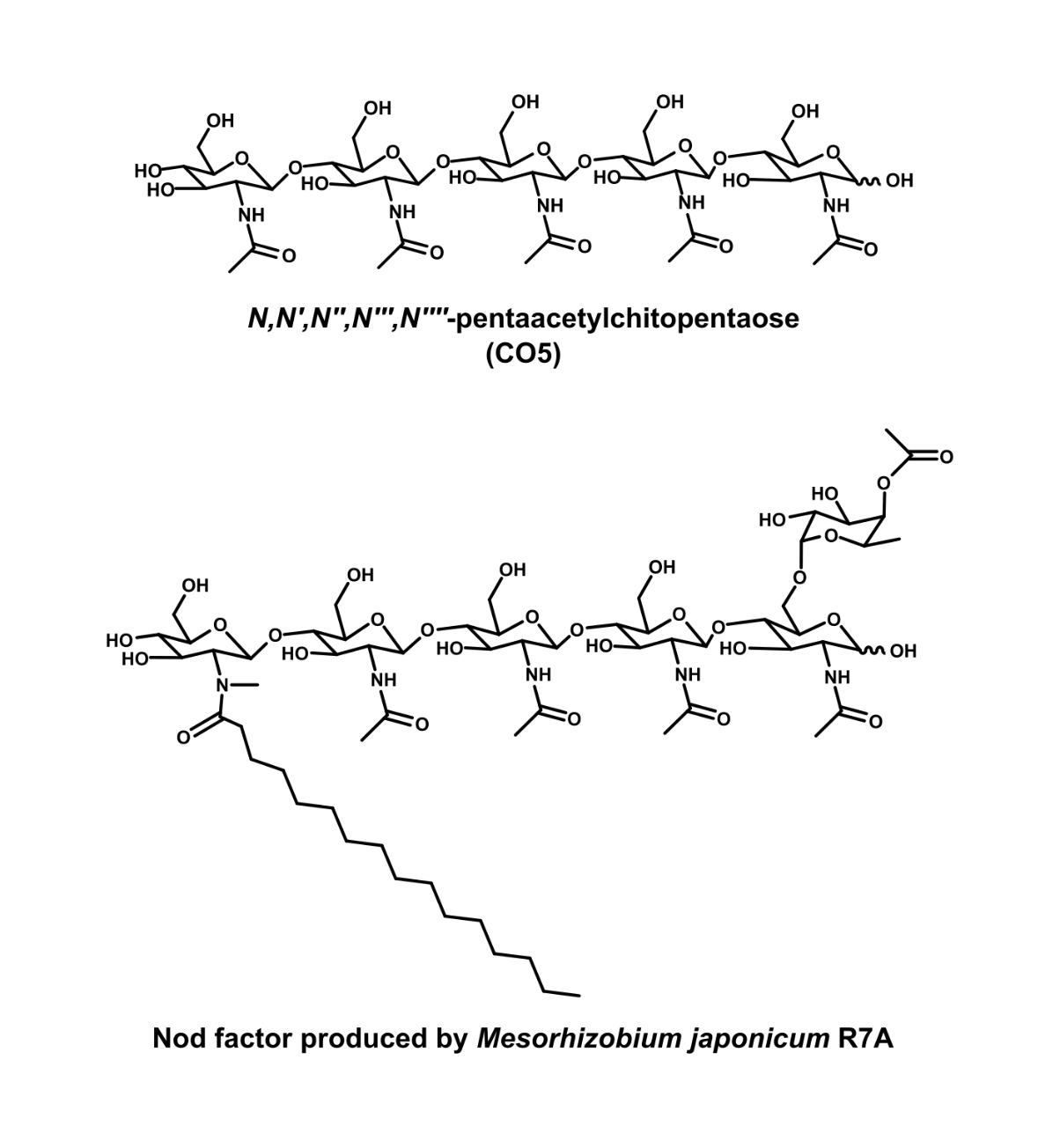

**Figure S1. Structures of CO5 and Nod factor.** In the top panel, the chemical structure of *N,N’,N’’,N’’’,N’’’’*-pentaacetylchitopentaose (i.e., CO5) is shown. In the bottom panel, an example of one Nod factor made by *M. japonicum* R7A (i.e., V(C16:0, NMe, AcFuc)) (9) is shown. In this molecule, the reducing terminal GlcNAc is decorated with an acetylated α-1,6-linked L-fucose moiety.

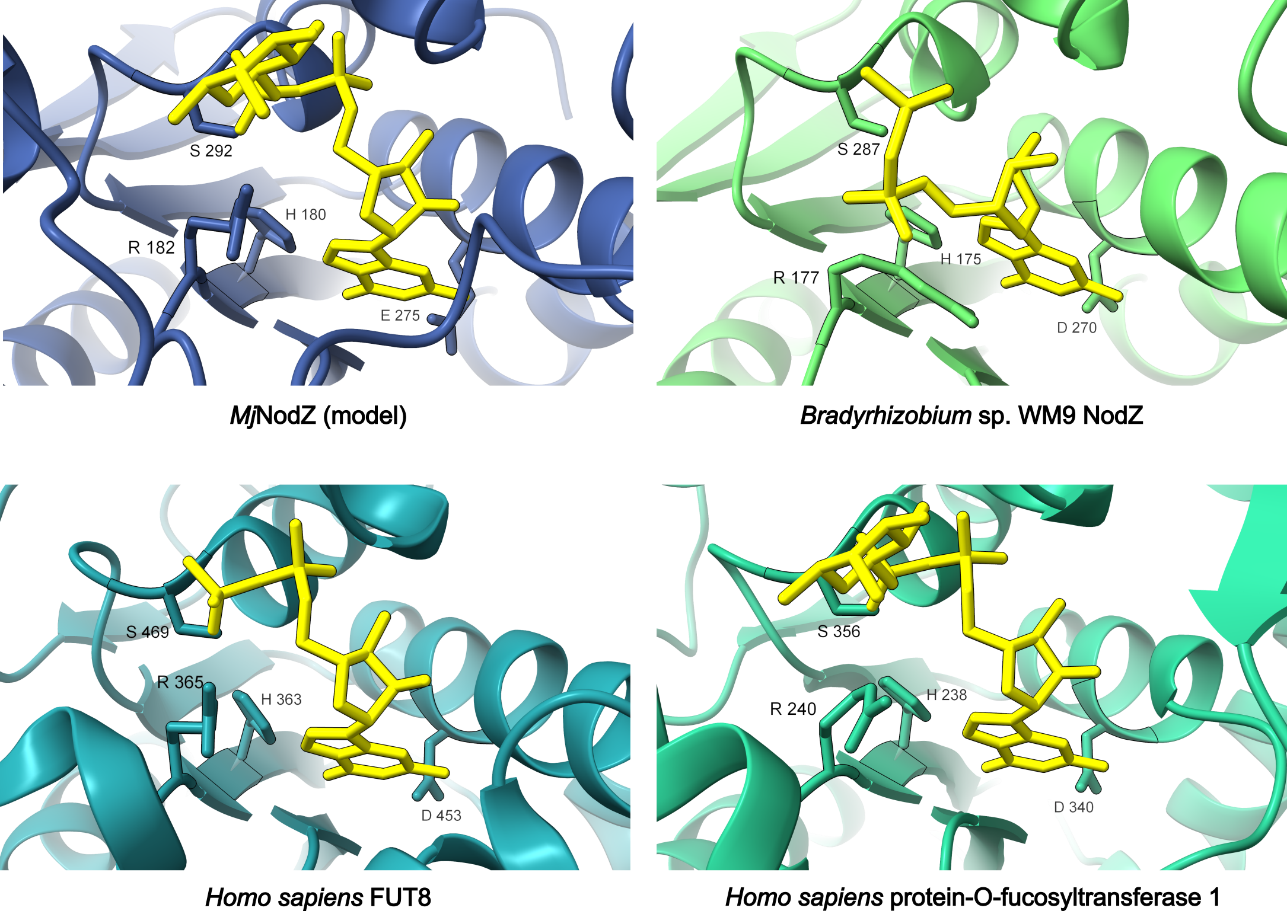

**Figure S2. Validation of the GDP-Fuc binding pocket in the Chai1 *Mj*NodZ model.** A comparison of the conserved GDP-binding sites between fucosyltransferases is shown. Protein structures were superimposed using ChimeraX matchmaker. GDP or GDP-Fuc is shown in yellow, depending on the available crystal structures. Four conserved GDP-binding residues reported from the crystal structure of NodZ (1) are labeled in each enzyme. In the top left is a snapshot taken from the Chai1 model of *Mj*NodZ (ipTM ~ 0.87), which is shown in full length in Figure 2 of the main text. The remaining three panels show crystal structures of various fucosyltransferases in complex with GDP/GDP-Fuc; *Bradyrhizobium* sp. WM9 NodZ (PDB: 3SIX), *Homo sapiens* FUT8 (PDB: 6X5H), and *Homo sapiens* protein-O-fucosyltransferase 1 (PDB: 5UXH).

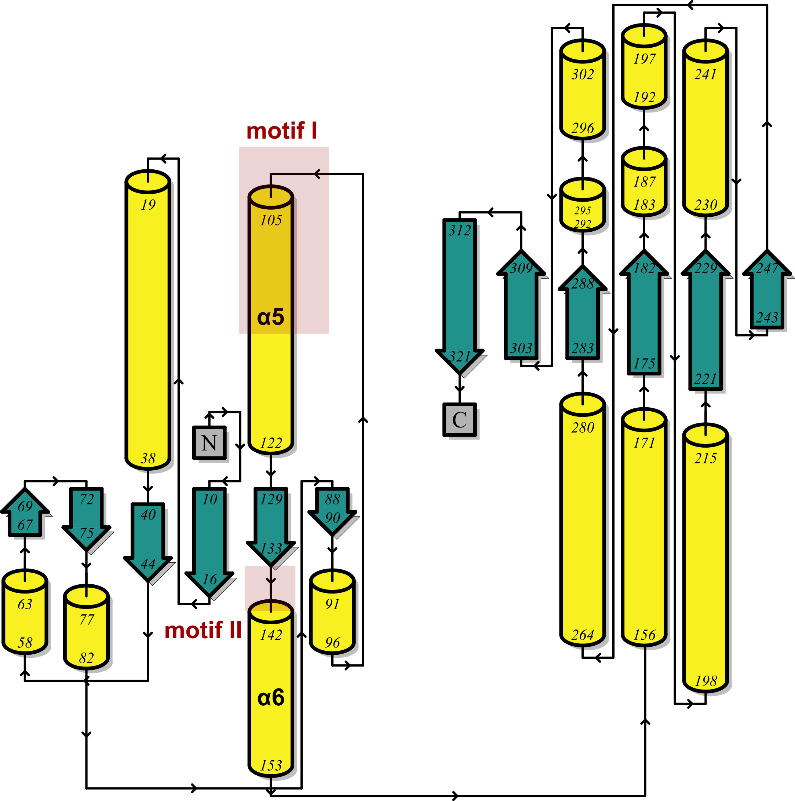

**Figure S3. Predicted secondary structure of *Mj*NodZ.** A cartoon of the secondary structural elements of *Mj*NodZ is shown. The secondary structure information was extracted from the AlphaFold2 model of *Mj*NodZ and submitted to the PDBsum web server to generate this cartoon. The α-helices are coloured in yellow and β-sheets are coloured in green. The regions representing motifs I and II are shown in red.

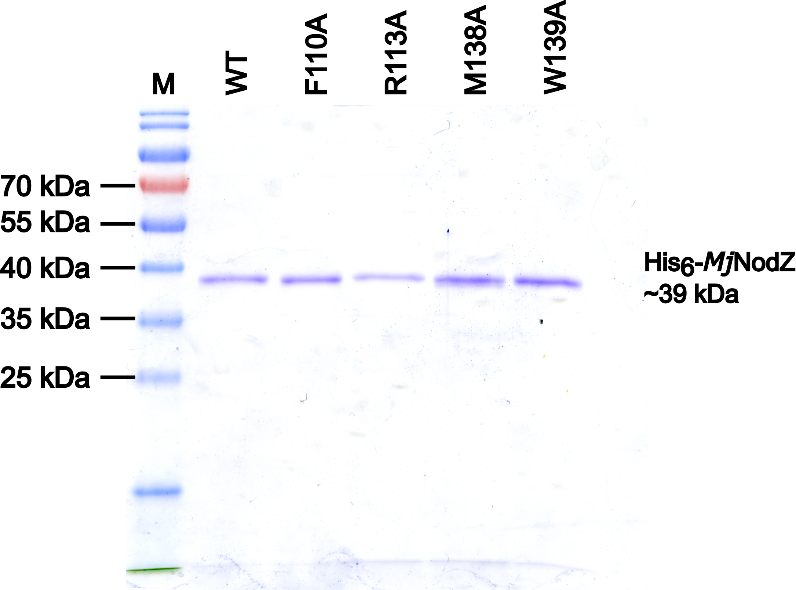

**Figure S4. SDS-PAGE analysis of purified *Mj*NodZ point mutants.** A 12% SDS-PAGE gel loaded with each recombinantly expressed and purified *Mj*NodZ protein mutant and stained with Coomassie Brilliant Blue R-250. The lanes are labelled according to the protein sample loaded, with WT (wildtype *Mj*NodZ) or with the corresponding mutation. An M is used to indicate the PageRuler Prestained Protein Ladder (Thermo Fisher). The expected mass of the protein is indicated on the right.

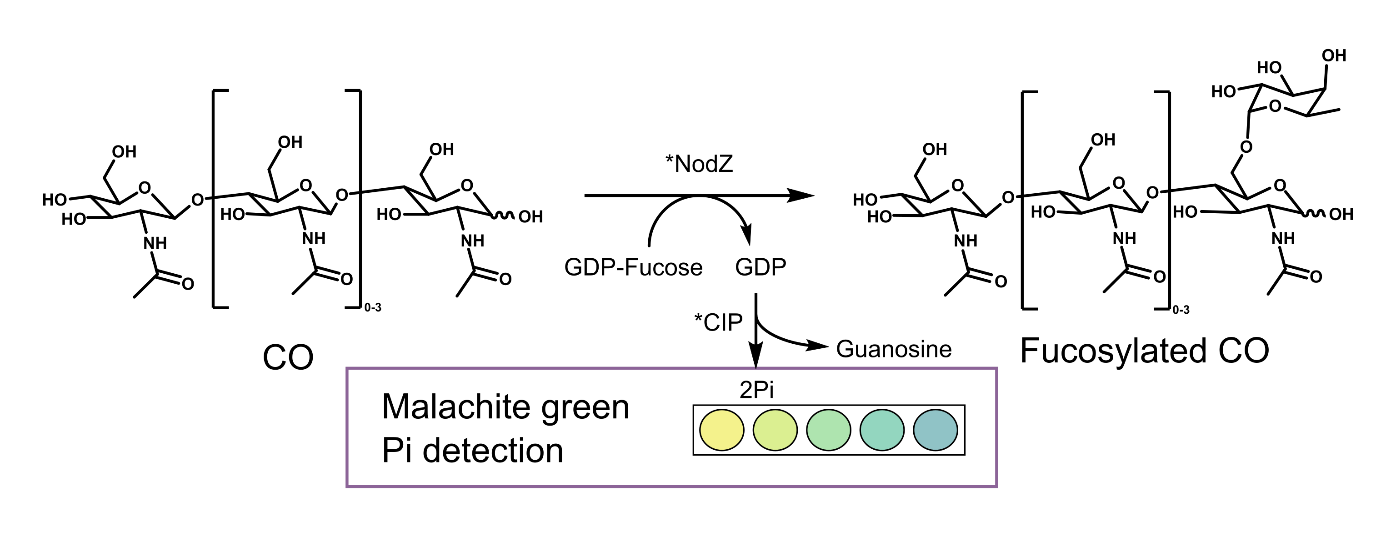

**Figure S5. Malachite green assay for measuring NodZ activity.** A simplified depiction of the malachite green kinetic assay is shown. A reaction scheme showing the NodZ-catalyzed fucosylation of a CO (chain length of two to five GlcNAc units), and the coupled breakdown of GDP by calf intestinal phosphatase (CIP). The produced inorganic phosphate (Pi) is reacted with malachite green reagent in a microwell plate, allowing its quantification via absorbance at 620 nm.

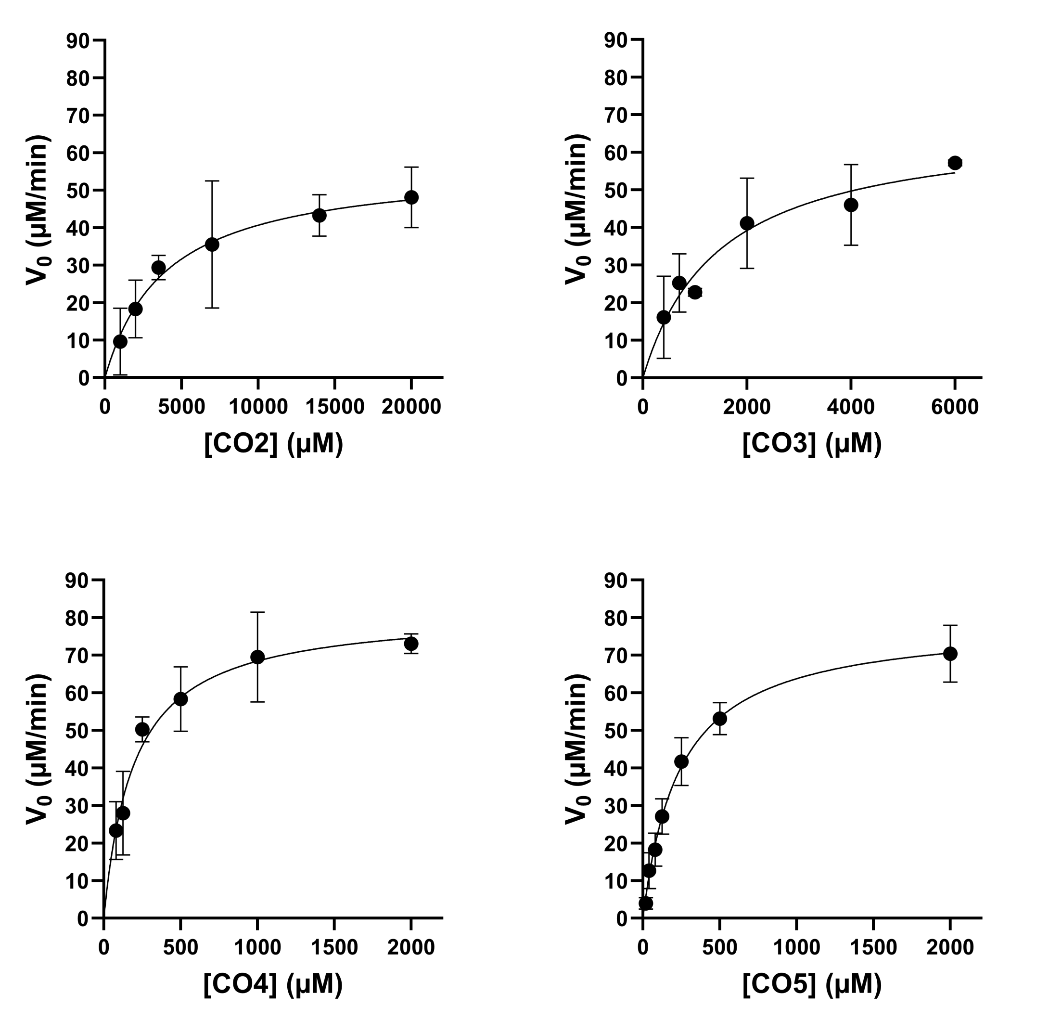

**Figure S6. Michaelis-Menten plots of *Mj*NodZ activity on different CO acceptors.** Plots of initial reaction velocity against substrate concentration are shown for wildtype *Mj*NodZ, with each plot representing one acceptor substrate (CO2, CO3, CO4, or CO5). Each point is the mean of at least three replicates and error bars represent standard deviation from the mean. The data were fitted to the Michaelis-Menten equation using GraphPad Prism with default settings. The parameters calculated from these fits are outlined in **Table 1** of the main text.

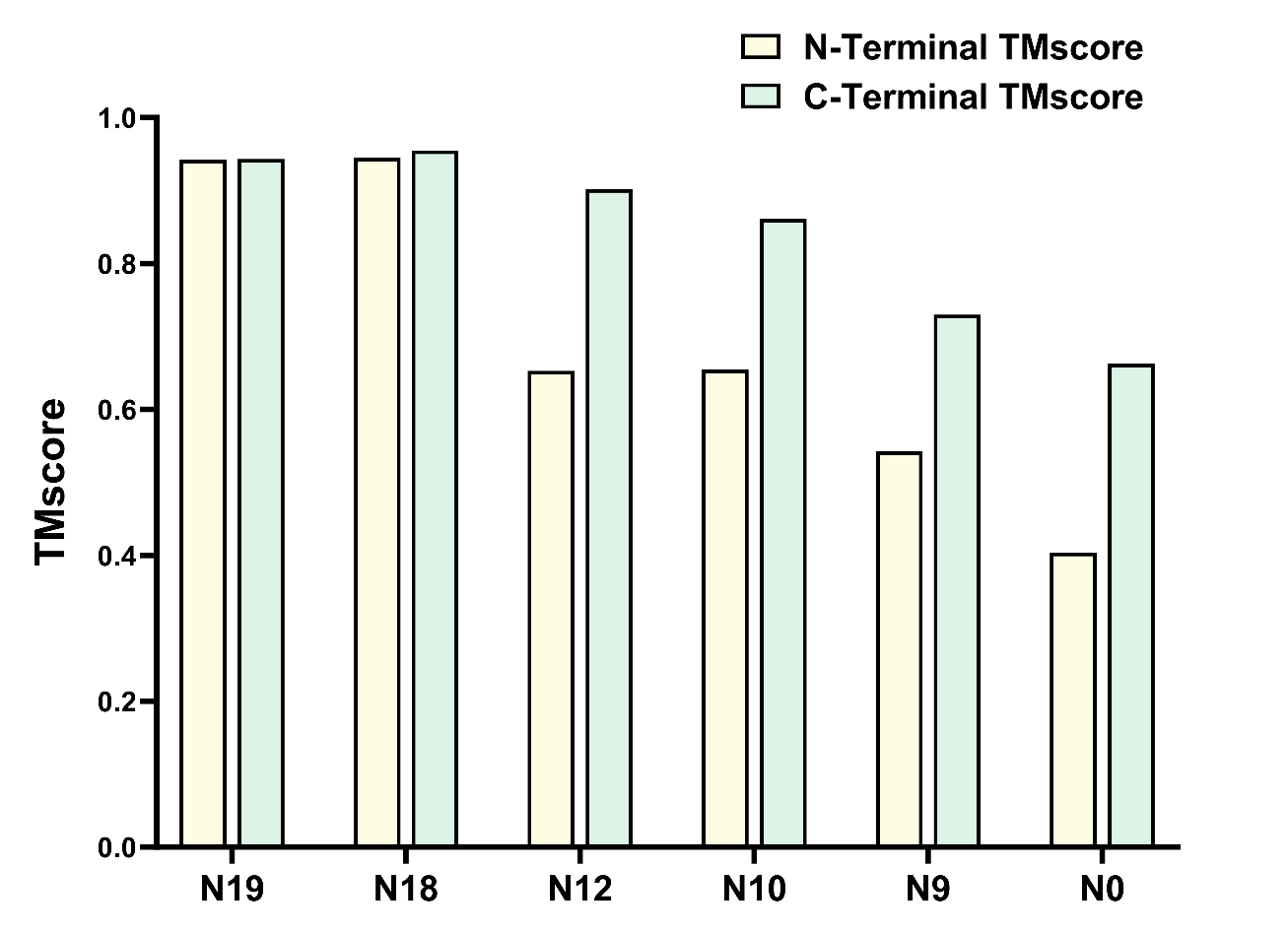

**Figure S7. TMscores of truncated models of ancestral proteins.** The ancestral NodZ proteins were modelled with AlphaFold2, truncated into their *N*- and *C*-terminal domains, then the truncated models were aligned with the model of *Bradyrhizobium* sp. WM9 NodZ with TM-align (13). This barplot shows the TMscore obtained for each *N*-terminal (yellow) or *C*-terminal (green) alignment.

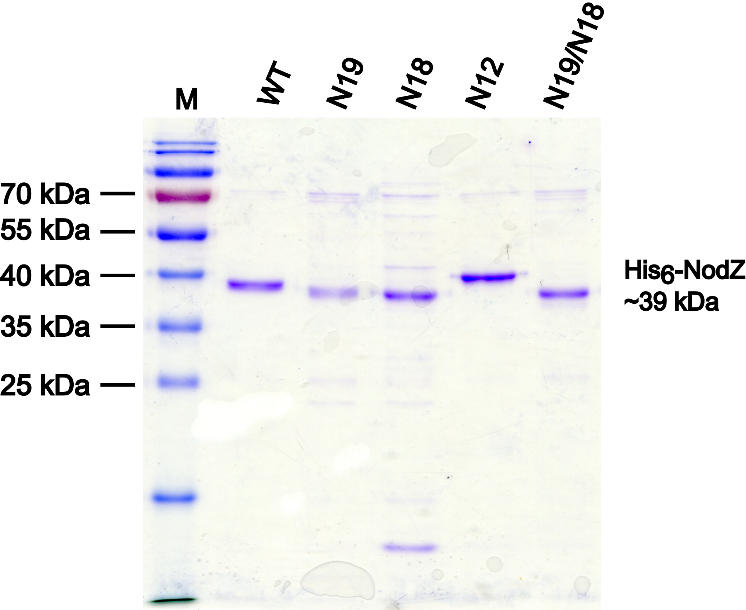

**Figure S8. SDS-PAGE analysis of purified ancestral NodZ proteins.** A 12% SDS-PAGE gel loaded with ancestral NodZ proteins/variants and stained with Coomassie Brilliant Blue R-250. The lanes are labelled according to the protein sample loaded, with WT (wildtype *Mj*NodZ) or with the ancestral protein. “N19/N18” refers to the chimeric fusion protein. An M is used to indicate the PageRuler Prestained Protein Ladder (Thermo Fisher). The approximate mass of the NodZ proteins is indicated on the right.

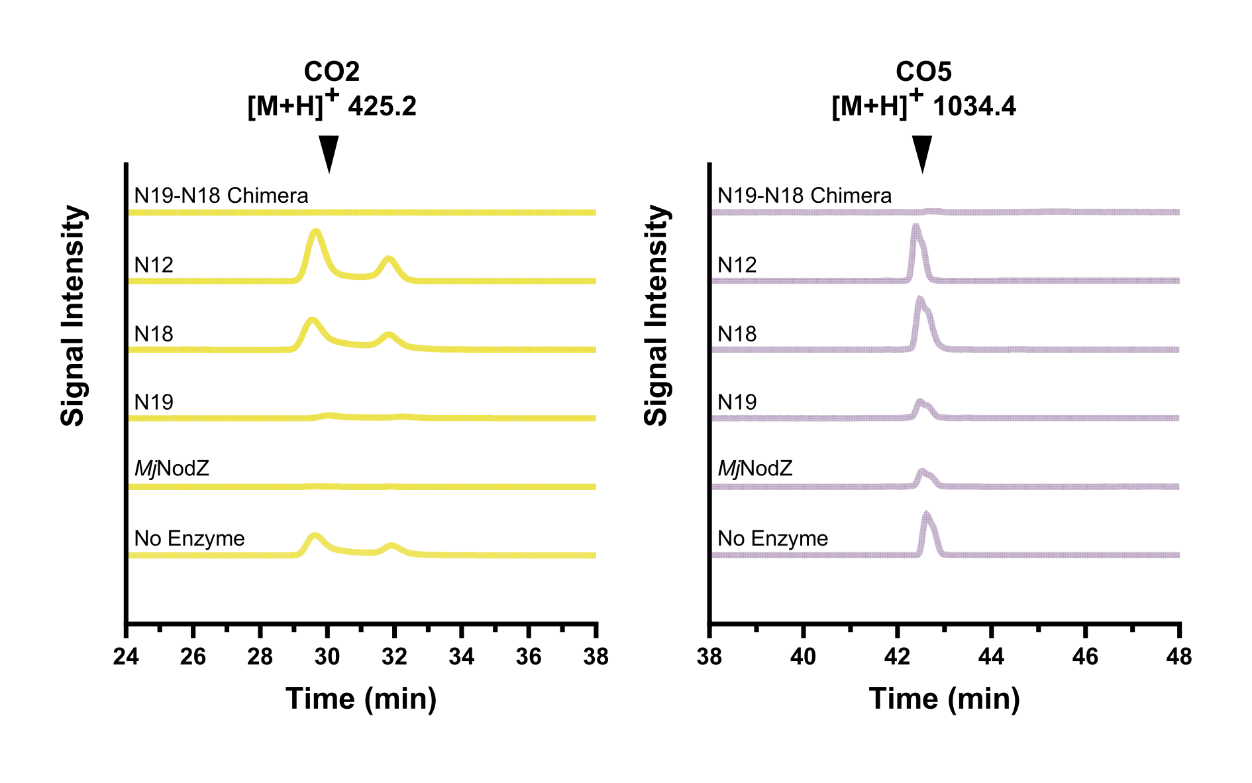

**Figure S9. Detection of CO2 and CO5 with HILIC-MS.** The detection of unmodified COs in reactions performed with ancestral NodZ enzymes and their mutants is shown. Chromatograms show the results of single-ion monitoring (SIM) for the [M+H]^+^ ion corresponding to the CO starting substrates.

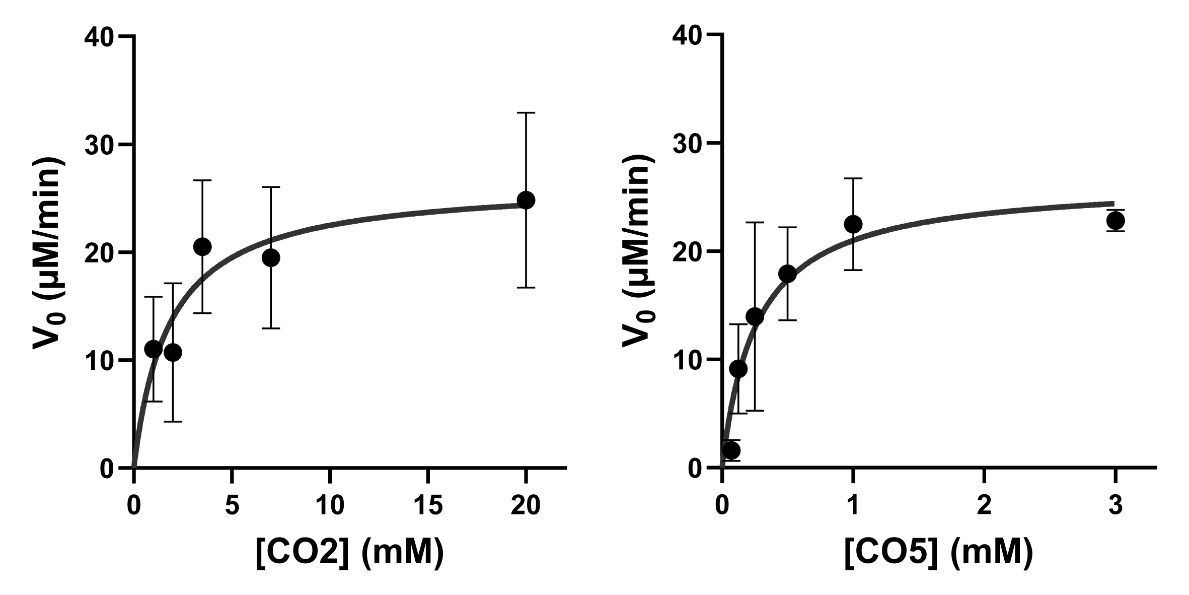

**Figure S10. Michaelis-Menten plots of N19 activity.** Plots of initial reaction velocity against substrate concentration are shown for the ancestral protein N19, with an acceptor substrate of either CO2 (left) or CO5 (right). Each point is the mean of at least three replicates and error bars represent standard deviation from the mean. The data were fitted to the Michaelis-Menten equation using GraphPad Prism with default settings.
